# A Curated Cell Life Imaging Dataset of Immune-enriched Pancreatic Cancer Organoids with Pre-trained AI Models

**DOI:** 10.1101/2024.02.12.580032

**Authors:** Ajinkya Kulkarni, Nathalia Ferreira, Riccardo Scodellaro, Frauke Alves

**Affiliations:** Translational Molecular Imaging, Max Planck Institute for Multidisciplinary Sciences, Hermann-Rein-Straße 3, 37075 Göttingen, Germany; Department of Haematology and Medical Oncology, University Medical Center Göttingen, Robert-Koch-Straße 40, 37075 Göttingen, Germany; Institute for Diagnostic and Interventional Radiology, University Medical Center Göttingen, Robert-Koch-Straße 40, 37075 Göttingen, Germany

**Author notes:** corresponding author(s): Frauke Alves. these authors contributed equally to this work (Ajinkya Kulkarni, Nathalia Ferreira).

## Abstract

Tumor organoids are three-dimensional in vitro models which can recapitulate the complex mutational landscape and tissue architecture observed in cancer patients, providing a realistic tumor microenvironment for testing novel therapies, including immunotherapies. A significant challenge in organoid research in oncology lies in developing efficient and reliable methods for segmenting organoid images, quantifying organoid growth, regression and response to treatments, as well as predicting the behavior of organoid systems. Up to now, a curated dataset of organoids co-cultured with immune cells is not available. To address this gap, we present a new public dataset, comprising both phase-contrast images of murine and patient-derived tumor organoids of one of the deadliest cancer types, the Pancreatic Ductal Adenocarcinoma, co-cultured with immune cells, and state-of-the-art algorithms for object detection and segmentation. Our dataset, *OrganoIDNetData*, encompassing 190 images with 33906 organoids, can be a potential common benchmark for different organoids segmentation protocols, moving beyond the current practice of training and testing these algorithms on isolated datasets.

## Background & Summary

Pancreatic Ductal Adenocarcinoma (PDAC) is the most prevalent neoplastic disease of the pancreas, accounting for over 90% of all pancreatic malignancies^1^. Characterized by its aggressive nature PDAC patients with metastasis have a median life expectancy <1 year. Such dismal prognosis can be traced back to late detection, tumor heterogeneity, intrinsic chemoresistance and failure of conventional therapeutic approaches. Moreover, the chemotherapeutic strategies as cornerstone of treatment often lead to increased side-effects and toxicity amongst the PDAC patients, accompanied with only a slight increase in survival rates^2^. Therefore, especially PDAC poses a formidable challenge in the field of oncology. Offering a distinct approach, immunotherapy has transformed cancer treatment strategies, demonstrating fewer side effects and reduced tumor resistance compared to conventional chemotherapy^3^. However, applying immunotherapeutic strategies to PDAC patients still faces obstacles. PDAC is an inherently immune-cold tumor and employs various strategies to counteract the effectiveness of immunotherapy, such as a dense stroma that i) hinders the infiltration of T-cells into the PDAC tumor microenvironment, ii) promotes chemoresistance and immune escape, and iii) produces cytokines that support the growth and survival of the tumor cells^4^. Additionally, the translation of immunotherapy into clinical applications is impeded by the inadequacy of effective pre-clinical models.

Although traditional animal models have been extensively utilized, they fall short in replicating the unique PDAC tumor microenvironment (TME), immune evasion mechanisms, and the distinctive genetic landscape of PDAC^5^. Acting as a bridge between two-dimensional cell lines and advanced patient-derived xenograft (PDX) models, organoids serve as a three-dimensional in vitro model capable of faithfully reproducing the complex mutational landscape observed in PDAC patients. Furthermore, patient-derived organoids (PDOs) further provide a platform to mimic the unique genetic characteristics of PDAC patients, advancing as a pre-clinical model that facilitates personalized treatment options in clinical practice. Organoids exhibit characteristics such as self-renewal and self-organization into mini-organ structures that closely resemble the original tissue architecture^6^. Despite the significant strides PDAC organoids bring to pre-clinical testing of novel therapies, they fall short in maintaining the immune system compartment of the PDAC-TME, posing a challenge for evaluating new immunotherapeutic strategies^7^.

To address the absence of the immune system compartment, researchers have been refining protocols to create an optimal environment that facilitates the co-culture of organoids with the immune system^8^. This endeavor has been instrumental in testing immunotherapies, including antibody-based immune checkpoints, leading to novel discoveries in the immunotherapeutic field^9^. The practical evaluation of treatment efficacy in organoids involves the use of Cell-Titer Glo as an endpoint assay, allowing the quantification of metabolic active cells and assessment of organoid viability^10^. Nevertheless, live cell imaging was discovered as a breakthrough technology to evaluate life response of organoids, thanks to its capability to monitor the real time reaction of organoids towards the therapy during every steps of the treatment^11–13^.

Up to now, several algorithms have emerged to analyze the response of organoids during live cell imaging to specific treatments based on physical measurements of organoid properties, such as mean area, organoid eccentricity and total organoid count. These algorithms enable the continuous assessment of organoid reactions over time without disrupting the organoid system^14–18^. Nevertheless, each algorithm is evaluated on its proper dataset, in optimal conditions, not allowing a valid comparison among the different performances. Moreover, all of these datasets report organoids in a relatively simple environment, where co-culture with immune cells has not been considered.

Here, we propose OrganoIDNetData, a public dataset of phase-contrast images of human and murine PDAC organoids, co-cultured with immune cells. This dataset enables to provide a curated common benchmark among different segmentation protocols, checking their performances in a more complex environment, incorporating pre-activated peripheral mononuclear cells (PBMCs) into the system. Together with the images, the dataset also provides the scripts to analyze the data with two different state-of-the-art algorithms for image segmentation and object detection, Cellpose^19^ and StarDist^20^.

## Methods

### Sample preparation

#### Human Cultures

All specimens of Pancreatic Ductal Adenocarcinoma (PDAC) tumors from human patients, used in the generation of PDAC Patient Derived Organoids (PDOs), were sourced during surgical intervention from individuals participating in the Molecular Pancreas Program (MolPAC) at the University Medical Center of Göttingen (UMG). This was done in accordance with the approval granted by local regulatory authorities (Ethics Committee of the UMG), as indicated by approval references 11/5/17, 22/8/21Ü, and 2/4/19. In the case of human Peripheral Blood Mononuclear Cells (PBMCs), peripheral blood was obtained from anonymous healthy donors with approval from the Ethics Committee of the UMG (ETHIC approval 29/07/23).

#### Murine Cultures

To generate murine PDAC tumors, a quantity of 5 *×* 10^5^ KPC cells was orthotopically implanted into the pancreatic head of C57BL/6 J mice, following previously established procedures^21^. Murine peripheral blood was collected from euthanized healthy C57BL/6J mice through cardiac puncture. All experiments were conducted with approval from the administration of Lower Saxony (approval number G20.3527) in accordance with current German laws governing animal experimentation.

#### Medium Preparation

- Digestion medium: Composition per 100 *ml* included 12 *mg* Collagenase type I (Sigma), 12 *mg* Dispase II (Sigma), and 1 *ml* of 10% FCS (Gibco) in 99 *ml* of DMEM (Gibco). Human Organoid Growth Medium (HOGM): 50 *µl* of HOGM consisted of 25 *µl* A83-01 (1*mM*, Tocris), 50 *µl* Human Epidermal Growth Factor (hEGF; 500 *µg/ml*, Invitrogen), 50 *µl* human Fibroblast Growth Factor-10 (hFGF-10; 100 *mg/ml*, Peprotech), 50 *µl* Gastrin I (100 *µM*, Sigma), 125 *µl* N-acetylcysteine (500 *mM*, Sigma), 500 *µl* Nicotinamide (1 *M*, Sigma), 1 *ml* B-27 supplement (50x, Gibco), 100 *µl* Primocin (50 *mg/ml*, InvivoGen), with additional components diluted in 19 *ml* of organoid splitting medium (1*x* Glutamax, 1*x* HEPES, 1 *ml* 1*x* Primocin, 30% Bovine Serum Albumin (BSA) diluted in Advanced DMEM/F12 medium (AdDMEM/F12, Gibco). For initial seeding, splitting, or thawing, 1:1000 Rho Kinase Inhibitor (Sigma) was added.
- Murine Organoid Growth Medium (MOGM): 50 *ml* of MOGM contained 5 *µl* murine Epidermal Growth Factor (mEGF; 500 *µg/ml*l, Invitrogen), 50 *µl* murine Fibroblast Growth Factor-10 (mFGF-10; 100 *µg/ml*, Peprotech), 5 *µl* Gastrin I (100 *µM*, Sigma), 125 *µl* N-acetylcysteine (500 *mM*, Sigma), 500 *µl* Nicotinamide (1 *M*, Sigma), 1 *ml* B-27 supplement (50x, Gibco), 5 *ml* R-spondin, and 5 *ml* of Noggin-conditioned media diluted in 38.3 *ml* of organoid splitting medium (1*x* Glutamax, 1*x* HEPES, 1% Penicillin-Streptomycin diluted in AdDMEM/F12). Similar to HOGM, 1:1000 Rho Kinase Inhibitor (Sigma) was added for initial seeding, splitting, or thawing.
- Wnt3a-, R-Spondin- and Noggin-conditioned media: Wnt3a-, R-Spondin-, and Noggin-conditioned media were prepared as detailed in previous literature^22,23^. Conditioning involved culturing specific cell lines (L-Wnt3A for Wnt3a, 293T-HA-Rspol-Fc for R-Spondin, and HEK293-mNoggin-Fc for Noggin) followed by collection, centrifugation, pooling, and filter sterilization before storage at -20°C.
- Organoid passaging medium (OPM): 500 *ml* of OPM included 5 *ml* 100x Glutamax, 5 *ml* 1 *M* HEPES, and 1% Penicillin-Streptomycin in 500 *ml* AdDMEM/F12.
- PBMCs culture medium: RPMI medium supplemented with 10% FCS, 50 *µM β* -mercaptoethanol, and 1% Penicillin-Streptomycin.

#### PDAC organoid establishment and culturing

PDAC organoids, obtained from either murine or human PDAC tumors, were generated following established organoid cell culture procedures outlined^24^. In summary, cells were isolated from freshly excised PDAC tumor samples and exposed to a digestion medium. Subsequently, the cells were suspended in 50 *µl* of Growth Factor Reduced (GFR) Matrigel (Corning) to form a dome within a preheated 24-well plate. Upon solidification of the dome, the respective organoid growth medium for human and murine PDAC organoids was applied to the top. Human and murine organoid formation was observed by live cell imaging (see the “Live cell imaging protocol” subsection for further information) after 3 days of culturing at 37°C in a humidified atmosphere with 5% *CO*_2_. The organoids were used for experimentation after undergoing a minimum of 3 passages.

#### PDAC organoids co-cultures with PBMCs

Following the 3-day formation of PDAC organoids, a co-culture with PBMCs was initiated. Simultaneously, human and murine PBMCs underwent pre-activation through culturing in PBMCs culture medium supplemented with ImmunoCultTM Human CD3/CD28 T cell Activator (StemCell Technologies) and suspension in LymphoGrow II medium (Cytogen), respectively. After an overnight pre-activation phase, co-cultures were established at a ratio of 10^4^ organoids to 10^5^ pre-activated PBMCs. The medium, comprising 50% MOGM or HOGM with 50% PBMC medium, was then added.

### Imaging protocol and dataset preparation

#### Live cell imaging protocol

The growth and viability of PDAC organoids and PBMCs organoid co-cultures were monitored with the live-cell imaging system IncucyteR S3 (Sartorius, Germany). The Incucyte S3 system from Sartorius proves to be a valuable asset for observing the time-lapse behavior of organoids in response to a particular treatment. This in-incubator microscope system offers label-free, automated acquisition of organoids within a physiologically stable environment favorable to organoid growth. 190 Phase-contrast fields of view of a mixture of sizes (1408 *×* 1040 pixels & 1280 *×* 852 pixels) were acquired every 4 hours up to 100 hours using the *Organoid* mode. The 4*x* objective provided a resolution of 2.82 *µm* per pixel.

#### Dataset Creation

The process of generating the proposed grayscale dataset used for training the model commenced with the extraction of the acquired field of views, as described by the Live cell imaging protocol. The dataset was then meticulously verified for integrity and completeness by using the “*count_image_names_and_check_masks*” function from the “*Prepare_Dataset_Modules*.*py*”. This function ensures that each field of view has a corresponding mask (generated by experts by manually annotating and segmenting the organoids) and identifies the quantity of fields of view belonging to human patients or murine cultures, by checking their prefixes “*human_*” and “*mouse_*”. Following this, the “*extract_patches*” function was employed to segment the larger RGB images and masks into smaller grayscale patches, respectively converted into 8-bit and 16-bit, and stored in TIFF format. The “*extract_patches*” function is also available as a separate package at the dedicated GitHub public repository^25^. The patching was guided by specific parameters, including the patch size of size 800 *×* 800 pixels, overlap percentage of 200 pixels in horizontal and vertical directions. A minimum threshold of 4 labels per patch was incorporated, ensuring that each patch contained sufficient data for effective training. These parameters can be personalized by the user by modifying the input values in the provided script.

Subsequent to patch creation, a representative subset of these patches was selected and visually inspected to confirm their suitability for training. This crucial step validated the quality and applicability of the patches. Following this, the dataset underwent a division into training, validation and testing sets based on a pre-determined ratio (75% for training and 20% for validation and 5% for testing). This split was carefully executed to ensure a balanced distribution of data. This is followed by the augmentation phase, using the augment function from “*Prepare_Dataset_Modules*.*py*” to each image in the training set. This function systematically applies a series of random transformations, such as flipping, rotation, translation, zooming, and adjustments in brightness and contrast—to introduce variability in the dataset. Each image was subjected to a specified number of augmentations (as determined by the “*n_augmentations*” variable, in this study fixed at 2), thereby significantly enriching the dataset with diverse representations of the original data.

The final stages of the process included comprehensive checks for data sanity and validation using the “*check_data_sanity*” and the “*validate_and_count_images*” functions. These functions ensure that the dataset maintains its structural integrity, adheres to the expected data types and value ranges (0-255 for the 8-bit format patches, and 0-65535 for the 16-bit format masks), and provide a transparent overview of its composition, including the count of human and mouse prefixed images and masks. Finally, the “*count_organoid_number_and_report*” function automatically counts and reports the number of organoids in the masks folder. It calculates both the total and average number of organoids across all the masks, ignoring the background.

#### Automated image segmentation protocols

To validate the dataset and demonstrate that it can be a potential challenging benchmark to compare different segmentation and object detection protocols, we analyzed it by training two state-of-the-art algorithms: Cellpose^19^ and StarDist^20^. Cellpose is a deep learning-based method for cellular segmentation of in microscopy images. Cellpose works by using a convolutional neural network (CNN) to identify and segment cells in images. Particularly suitable for fluorescence and phase-contrast images, Cellpose is characterized by a great versatility, since the CNN is trained on a dataset including images of cells of different types, sizes, and conditions. Characterized by a simpler architecture than Cellpose, StarDist is a deep-learning method primarily designed for segmenting cells in vitro and in vivo. It also relies on CNNs, and uses a star-convex polygon approach to segment objects by defining convex polygons around them. StarDist is particularly designed for addressing challenges in segmenting elongated and branched structures in microscopy images. Both the algorithms were trained for 1000 epochs and the weights were saved for future inference. Additionally, all other parameters which are used to train both models are also available in the GitHub repository^26^ as a part of the source codes.

## Data Records

The OrganoIDNetData dataset consists of systematically organized grayscale microscopy images, in TIFF format, and corresponding segmented mask files, aimed at supporting organoid research. The data and source codes are available on our dedicated GitHub repository^26^. For a clear understanding of the file structure, we provided a diagram in Figure 1, which depicts the three subfolders contained in the OrganoIDNetData main directory.

**Figure 1.**
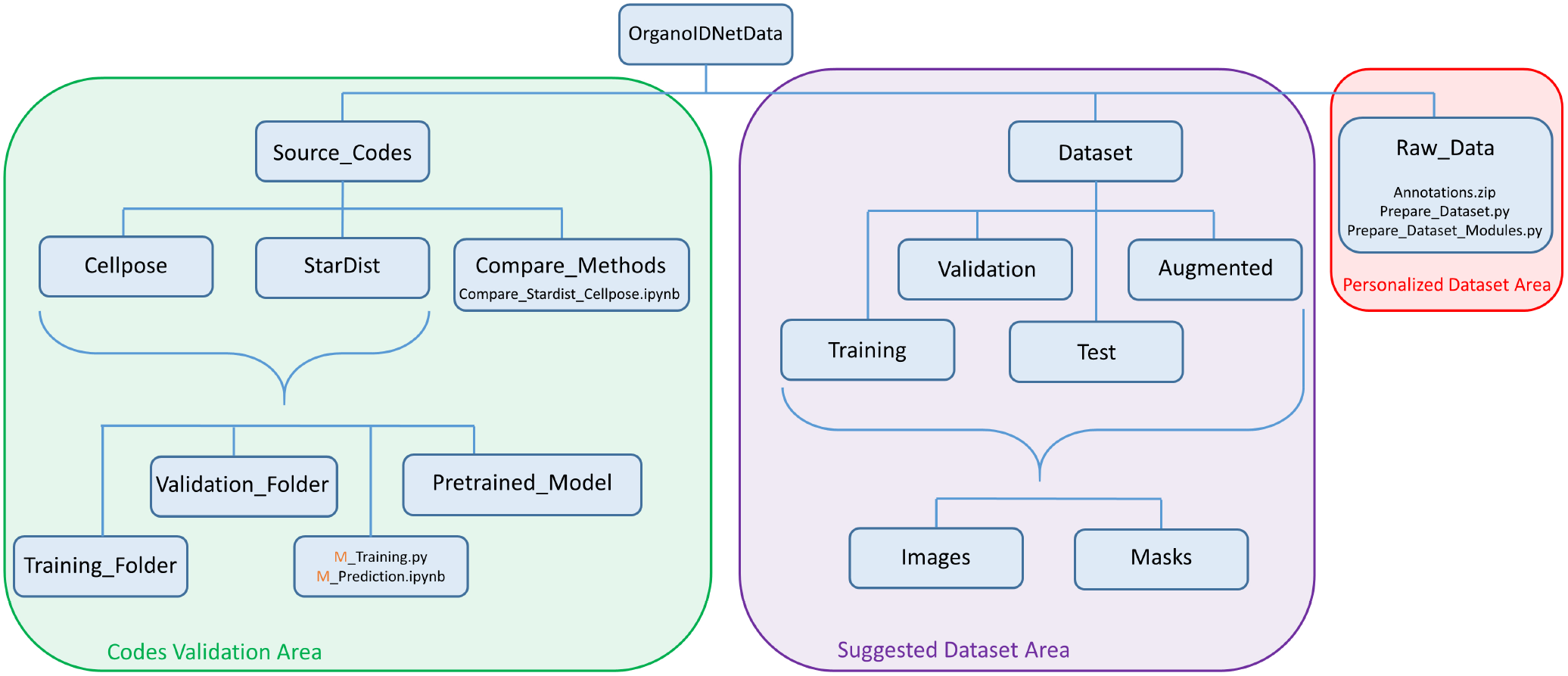
OrganoIDNetData file structure. Three main subfolders are contained in the OrganoIDNetData repository, recognizing three different areas of the directory. The “*Source_Codes*” subfolder stores all the python codes needed to reproduce the technical validation performed by using Cellpose and StarDist algorithms. The “*Raw_Data*” subfolder houses all the raw field of views, experimentally acquired to create the OrganoIDNet dataset, together with the python scripts that the user can modify and create its personalised version of the dataset. The “*Data*” subfolder stores the complete dataset presented in this article, organised in training, augmented, validation and test datasets.

The “*Source_Codes*” subfolder houses the Python codes used to perform the model training and conduct the technical validations of the manuscript. For both algorithms, Cellpose and StarDist, users can replicate the results by running these Python scripts. The “*M_Training*.*py*” script provides the training steps for each method, where M describes the approach (Cellpose or StarDist), while “*M_Prediction*.*ipynb*” can be used to analyze test data, once the method is trained. The data required for this process is readily available in the “*Training_Folder*” and “*Validation_Folder*” subfolders of the corresponding method. Alternatively, users can upload their own images to these subfolders and run the Cellpose and StarDist codes using their own data. However, both architectures, already trained with the proposed dataset, are stored in the “*Pretrained_Model*” subfolder. Additionally, the “*Compare_Methods*” folder houses the scripts necessary to obtain the quantitative metrics able to provide a comparison among different training methods. At the moment, the scripts are limited to the comparison of Cellpose and StarDist.

The “*Raw_Data*” folder houses a compressed archive, called Annotations.zip containing all the raw fields of view used to create the dataset. 94 fields of view belong to 4 different human cultures, while the remaining 86 belong to 2 different murine cultures. Moreover, this subfolder stores two Python scripts: “*Prepare_Dataset*.*py*” and “*Prepare_Dataset_Modules*.*py*”. These scripts are pre-configured to process the raw field of views and generate the dataset presented in this article. The scripts are commented to enhance the versatility of the dataset by letting the user create its personalised version of the dataset. However, the code can be modified to create new datasets with different image dimensions.

The “*Dataset*” folder also stores the complete dataset presented in this article, obtained by all the raw field of views. The images are organized into distinct subfolders. The “*Training*” subfolder contains 713 images (364 from human cultures, 349 from murine cultures) of 800 *×* 800 pixels, which are used to train the network. Conversely, the “*Validation*” subfolder holds 179 (98 from human cultures, 81 from murine cultures) images, which are used to optimize the network’s performance and parameters during the training. The “*Augmented*” subfolder contains 1426 augmented images generated by applying data augmentation on the training set. These augmented images were added to the standard training set to expose the network to a wider variety of image representations and used during the technical validations presented in this article. The “*Test*” subfolder contains 10 entire fields of view (5 from human cultures, 5 from murine cultures), not splitted in patches, that can be used to evaluate the network’s performance on unseen data. Within each of these subfolders, images are further organized into two subfolders: the “*Images*” subfolder contains the actual images comprising our dataset, while the “*Masks*” subfolder stores the corresponding masks for each image, which were obtained by manual annotation.

Both images and masks are saved in TIFF format and named using a standard convention that encodes their characteristics. The file name structure follows the pattern “*type_X_Wh_patchY_augZ*.*tif* “, where type *X* specifies the sample origin, either Human or Mouse. *X* is a unique identifier for the sample, *W* recognizes the time step of the measurement, expressed in hours, and patch identifies a distinct 800 *×* 800 pixels Y patch extracted from the same sample. The presence of the “*_augZ*” string is optional and indicates that the image was obtained through data augmentation, where the *Z* number distinguishes augmented data generated from the same image with different augmentation parameters. The mask filename follows the same pattern as the corresponding image filename, but with the final string “*_mask*” appended. For instance, the file “*Human_1_016h_patch4*.*tif* “recognizes the fourth patch extracted by the field of view acquired for the first human sample after 16 hours from the beginning of the live cell imaging acquisition. Conversely, “*Human_1_016h_patch4_aug2*.*tif*” is associated to the second augmented image generated by the original one, while their corresponding masks are “*Human_1_016h_patch4_mask*.*tif*” and “*Human_1_016h_patch4_aug2_mask*.*tif* “. When an entire field of view is provided, as in the case of the “*Raw_Data*” and “*Test*” subfolders, the string “*_patchY*” is not present in the name of the files. Moreover, each field of view we propose as part of the test dataset, and present in the “*Test*” subfolder, is characterized by “*Test_*” at the beginning of their name, following the structure “*Test_type_X_Wh*.*tif* “. Figure 2 displays two exemplary images (and their respective masks) from the dataset: one featuring organoids derived from a human patient and the other illustrating organoids from a murine sample. Some immune cells present in these two images are highlighted by manual annotation.

**Figure 2.**
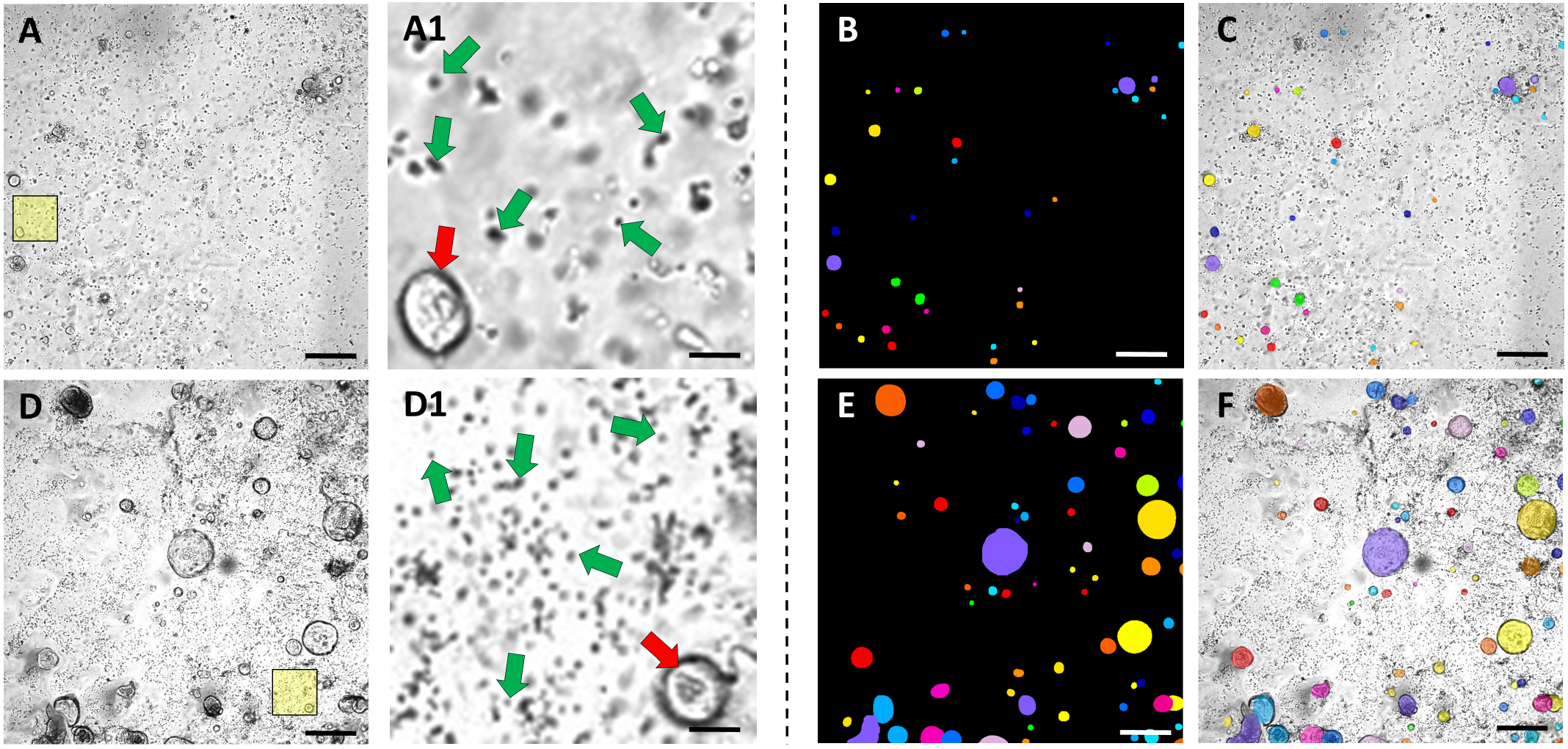
Representative images and corresponding masks of the OrganoIDNetData dataset. (A) and (D) report two dataset images, “*Human_1_024h_patch3*.*tif*” and “*Mouse_1_042h_patch0*.*tif*”, respectively. The two yellow squares highlight the zoomed-in regions of interest shown in (A1) and (D1). In each region of interest, a single organoid is indicated by a red arrow, while five immune cells, randomly selected, are highlighted by green arrows. (B) and (E), whose files are named “*Human_1_024h_patch3_mask*.*tif*” and “*Mouse_1_042h_patch0_mask*.*tif*”, show the corresponding manual segmentations for images (A) and (D), respectively. (C) and (F) depict the overlay between the original images of the dataset and the masks to better highlight the quality of the annotation. Scale bars: 115 *µm* for images from (A) to (F), 15 *µm* for images (A1) and (D1).

## Technical Validation

Here, we present some proof-of-concept demonstrations showcasing the functionalities available to users with OrganoIDNetData. Given our proposition of the dataset as a potential common and standardized benchmark for various segmentation protocols, we conducted a comparative analysis of the outcomes obtained by applying StarDist and Cellpose. Furthermore, we analyzed a subset of the time-series live cell images present in the dataset, as it represents a potential feature that can be leveraged by the user. All analysis presented here was performed on a desktop workstation with an AMD Ryzen 9 5900X 12-Core Processor with 128 GB RAM, and an NVIDIA GeForce RTX 3080 GPU with dedicated 12 GB RAM.

### StarDist and Cellpose-based segmentation

Firstly, we used the corresponding subsets of the OrganoIDNetData dataset to train and test StarDist and Cellpose algorithms. For each image of the test dataset, both algorithms provided a segmentation mask, which was compared with the ground truth mask (i.e. the manual annotation) by computing 3 different quantitative metrics.

Intersection over Union (IoU) measures the degree of overlap between the algorithm’s predictions and the ground truth, with a higher IoU indicating better segmentation quality. The IoU is given by the formula: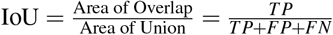, where *TP* are the true positives, *FP* are the false positives, and *FN* are the false negatives. The Dice Coefficient quantifies the similarity between algorithmic predictions and ground truth, with values closer to 1 indicating more accurate segmentation. It is calculated as Dice 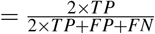. Finally, Pixel Accuracy assesses the accuracy of pixel-level segmentation, reflecting the reliability of the algorithm in segmenting individual pixels. It is defined as Pixel Accuracy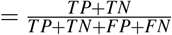, where *TN* represents the true negatives. These metrics offer distinct insights into the performance of organoid segmentation algorithms.

Regarding the quantitative assessment of organoid segmentation algorithms shown in Table 1, both StarDist and Cellpose demonstrated high performance across human and murine tissue images, as particularly evidenced by the Pixel Accuracy parameter. Despite the good results characterizing this metric, the lower values of Dice coefficients and Intersection over Union (IoU) suggest a modest divergence in the precise demarcation of organoid contours. This discrepancy underscores a potential avenue for algorithmic refinement, particularly in applications where boundary precision is paramount.

**Table 1.** Quantitative assessment of Organoid Segmentation Algorithms. This table presents the evaluation metrics for the performance of two organoid segmentation algorithms, StarDist and Cellpose, compared to ground truth masks. The metrics include Intersection over Union (IoU), Dice Coefficient, and Pixel Accuracy, calculated for each algorithm. The results showcase the accuracy and reliability of both algorithms in segmenting organoids from microscopy images at various time points. Higher values indicate better segmentation performance.

| Image | StarDist<br>IoU | StarDist Dice<br>Coefficient | StarDist Pixel<br>Accuracy | Cellpose<br>IoU | Cellpose Dice<br>Coefficient | Cellpose Pixel<br>Accuracy |
| --- | --- | --- | --- | --- | --- | --- |
| *Test_Human_1_004h.tif* | 0.660 | 0.795 | 0.991 | 0.736 | 0.848 | 0.994 |
| *Test_Human_1_008h.tif* | 0.684 | 0.812 | 0.991 | 0.704 | 0.826 | 0.992 |
| *Test_Human_1_012h.tif* | 0.714 | 0.833 | 0.993 | 0.742 | 0.852 | 0.993 |
| *Test_Human_1_016h.tif* | 0.665 | 0.799 | 0.991 | 0.678 | 0.808 | 0.992 |
| *Test_Human_1_020h.tif* | 0.659 | 0.795 | 0.992 | 0.727 | 0.842 | 0.994 |
| *Test_Mouse_2_000h.tif* | 0.647 | 0.786 | 0.914 | 0.689 | 0.816 | 0.925 |
| *Test_Mouse_2_002h.tif* | 0.662 | 0.796 | 0.907 | 0.675 | 0.806 | 0.911 |
| *Test_Mouse_2_004h.tif* | 0.667 | 0.800 | 0.892 | 0.692 | 0.818 | 0.901 |
| *Test_Mouse_2_006h.tif* | 0.672 | 0.804 | 0.891 | 0.691 | 0.817 | 0.899 |
| *Test_Mouse_2_008h.tif* | 0.668 | 0.801 | 0.891 | 0.674 | 0.805 | 0.894 |

In particular, concerning the human tissue images (‘*Test_Human1_004h*.*tif* ‘to’*Test_Human1_020h*.*tif*’), both algorithms showed comparable performance, though Cellpose slightly outperformed StarDist, notably in ‘*Test_Human1_004h*.*tif* ‘and’*Test_Human1_020h*.*tif* ‘with IoU scores of 0.736 and 0.727 and Pixel Accuracy values of 0.993 and 0.994 respectively. In murine tissue images (‘*Test_Mouse2_000h*.*tif* ‘to’*Test_Mouse2_008h*.*tif*’), a noticeable performance decrement was observed for both algorithms. Cellpose generally maintained a higher performance level, especially in terms of IoU and Dice Coefficients, for example, as seen in ‘*Test_Mouse2_008h*.*tif*’with an IoU of 0.674. Pixel Accuracy fared high and relatively consistent across all samples for both algorithms, albeit slightly lower in murine organoids. These findings indicate that while both StarDist and Cellpose are effective for organoid segmentation in microscopy images, Cellpose demonstrates a slight advantage, particularly in more challenging samples such as murine tissues, owing to a high density of organoids present, compared to the human samples (where organoids are fewer and sparsely distributed).

### Live cell imaging analysis

OrganoIDNetData can be also used to test the performance of novel algorithms in the analysis of time-series. By taking advantage of both StarDist and Cellpose, we analyzed two different time-series provided in the “*Test*” subfolder from the dataset, belonging to a human patient, specifically to a human sample (Human_1), and to a murine sample (Mouse_2). The overall results are provided in Figure 3. The murine and human samples of the test dataset are characterized by a very different density of organoids, whose number spans roughly 1700 and 70 organoids per image, respectively. In both scenarios, StarDist and Cellpose reflect the trend characterizing the ground truth over time. Nevertheless, Cellpose typically identifies more organoids in the human samples compared to StarDist.

**Figure 3.**
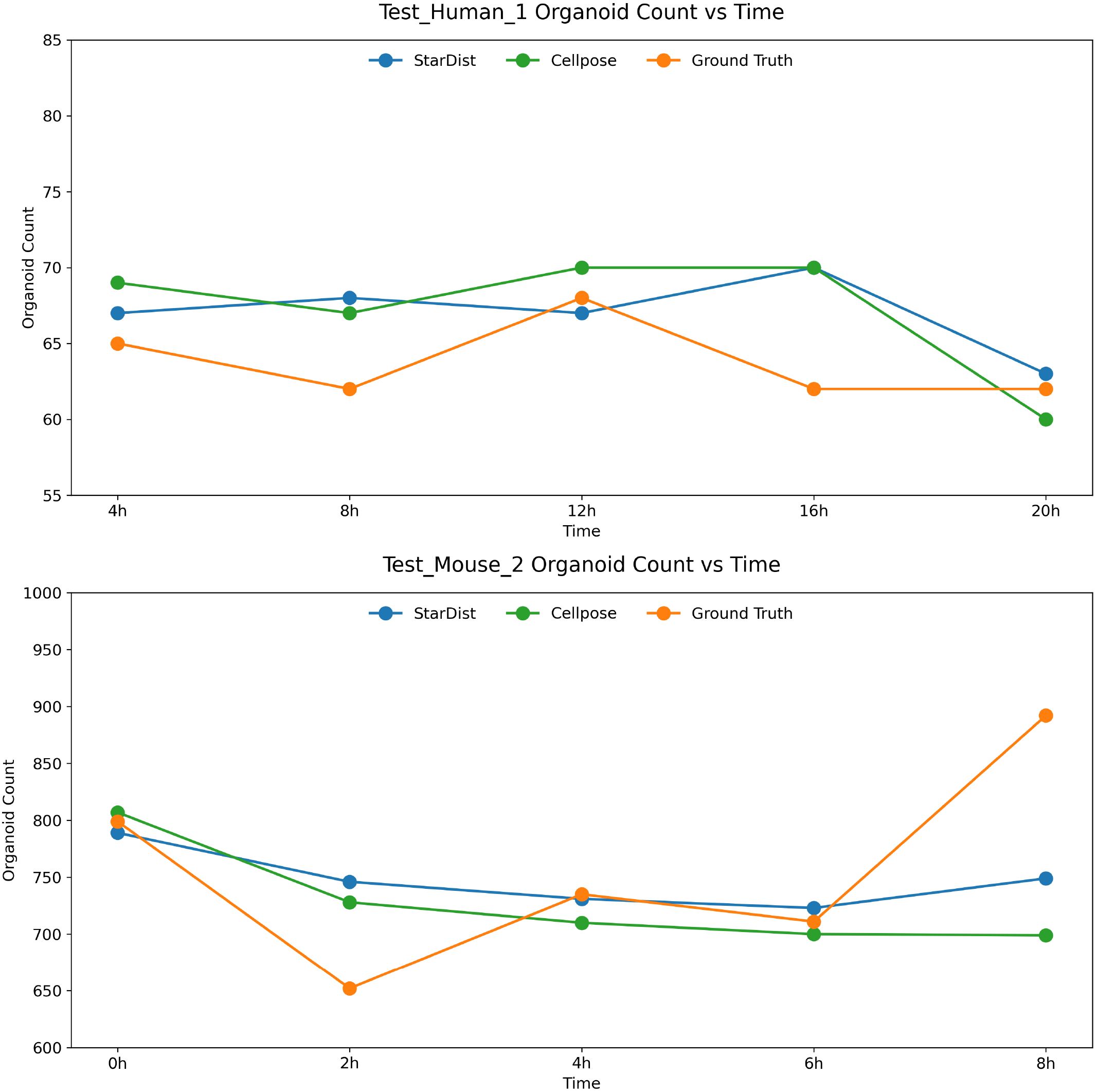
Comparative analysis between StarDist and Cellpose of two OrganoIDNetData time-series. (A) shows the organoid count obtained by Cellpose (green), and StarDist (blue), compared with the manual annotation outcome (orange) obtained during the analysis of the time-series involving the phase-contrast images acquired from the first human patient (recognized as “*Test_Human_1*” in the dataset). (B) shows the results obtained by performing the same analysis on the time-series involving the phase-contrast images acquired for a murine sample, specifically recognized in the dataset as “*Test_Mouse_2*”.

## Usage Notes

In our knowledge, OrganoIDNetData represents the first imaging dataset involving organoids co-cultured with immune cells. It is distinguished by meticulous manual annotations for every field of view. Its dimension, in terms of number of images, variety and number of organoids, is similar or higher than other datasets used to validate novel segmentation protocols^14,16,18^. The substantial variability in organoid density across the dataset images makes it a particularly challenging benchmark for the evaluation of segmentation protocols. The manual annotation, executed by experts, suggests an opportunity for enrichment with annotations contributed by more scientists. This collaborative effort would enhance the ground truth quality, mitigating potential subjectivity arising from a single expert. Up to now, the OrganoIDNetData dataset, whose images are consistently acquired within the same laboratory under similar conditions, is limited to murine and human PDAC samples, co-cultured with immune cells, and focuses on providing a challenging and common benchmark for object detection and segmentation AI-driven algorithms, as the proposed Cellpose and StarDist.

Similarly to initiatives undertaken for tumor samples^27^, OrganoIDNetData marks an initial stride towards establishing a more complex collaborative dataset. Such a dataset will provide an interesting data collection for many different AI algorithms, from multiclass classification to unsupervised learning-based protocols. Indeed, it would encompass the population’s diversity, featuring both images and numerical data, spanning a broad spectrum of organoids and their respective treatments. The provided scripts, leveraging Cellpose and StarDist-based analysis are properly commented to facilitate user customization. Moreover, each code is designed for compatibility with other datasets, provided they adhere to the same organizational structure as the proposed one.

## Code availability

The codes used in this article are available for free in the dedicated GitHub public repositories https://github.com/ajinkya-kulkarni/PyOrganoIDNet and https://github.com/ajinkya-kulkarni/PyBlendPatches, under the GNU General Public License v3.0. The pre-trained model weights and the dataset presented are available at 10.5281/zenodo.10643409, and will be freely accessible after publication in a peer reviewed journal.

## Acknowledgements

This project was in part funded by the Ministry for Science and Culture of Lower Saxony as part of the project “Agile, bio-inspired architectures” (ABA) under the grant agreement No ZN3822, and has received funding from the European Union’s Horizon 2020 research and innovation program under the Marie Skłodowska-Curie grant agreement No 861190– PAVE. The authors would like to thank Prof. Philipp Ströbel and Jennifer Appelhans, Institute of Pathology at the UMG for their assistance in cultivating PDAC organoids. We also like to thank Prof. Tobias J. Legler, Central Department of Transfusion Medicine at UMG for providing human immune cells. We also like to to acknowledge Daniele Ferrari for furnishing the murine KPC-PDAC tumor samples. We thank Bärbel Heidrich and Regine Kruse for their technical assistance. We express our gratitude to Sartorius for their technical assistance.

## Author contributions statement

A.K. wrote the Cellpose and StarDist codes and performed the technical validations, N.F. performed the experiments on organoids and acquired the raw images of the dataset. R.S. created the data record structure. F.A. supervised the project and provided financial support. All authors wrote and reviewed the manuscript.

